# Quantitative cytoarchitectural phenotyping of deparaffinized human brain tissues

**DOI:** 10.1101/2024.09.10.612232

**Authors:** Danila Di Meo, Michele Sorelli, Josephine Ramazzotti, Franco Cheli, Samuel Bradley, Laura Perego, Beatrice Lorenzon, Giacomo Mazzamuto, Aron Emmi, Andrea Porzionato, Raffaele De Caro, Rita Garbelli, Dalila Biancheri, Cristiana Pelorosso, Valerio Conti, Renzo Guerrini, Francesco S. Pavone, Irene Costantini

## Abstract

Advanced 3D imaging techniques and image segmentation and classification methods can profoundly transform biomedical research by offering deep insights into the cytoarchitecture of the human brain in relation to pathological conditions. Here, we propose a comprehensive pipeline for performing 3D imaging and automated quantitative cellular phenotyping on Formalin-Fixed Paraffin-Embedded (FFPE) human brain specimens, a valuable yet underutilized resource. We exploited the versatility of our method by applying it to different human specimens from both adult and pediatric, normal and abnormal brain regions. Quantitative data on neuronal volume, ellipticity, local density, and spatial clustering level were obtained from a machine learning-based analysis of the 3D cytoarchitectural organization of cells identified by different molecular markers in two subjects with malformations of cortical development (MCD). This approach will grant access to a wide range of physiological and pathological paraffin-embedded clinical specimens, allowing for volumetric imaging and quantitative analysis of human brain samples at cellular resolution. Possible genotype-phenotype correlations can be unveiled, providing new insights into the pathogenesis of various brain diseases and enlarging treatment opportunities.

## Introduction

In the realm of clinical and biomedical research, the advent of sophisticated 3D imaging methodologies heralds a profound evolution, offering unprecedented insights into the complexity of human brain architecture and pathology. Standard histopathological analysis, mainly performed on Formalin-Fixed Paraffin-Embedded (FFPE) tissue, has long been the cornerstone of clinical research and diagnostics [3, 24]. However, while providing valuable insights into tissue morphology and composition, histological techniques are inherently limited to two-dimensional (2D) representations, leading to a loss of spatial context and a failure to capture the intricate complexities of biological structures. Conversely, volumetric imaging analysis offers a comprehensive understanding of the spatial organization of human brain areas enabling researchers to capture the full complexity of biological structures and correlate it to their function [59]. Histopathological analyses are routinely performed on ultra-thin slices (<10-μm-thick) of FFPE specimens. As a result, only a fraction of the FFPE tissue block is utilized, while the remaining is often archived and stored for extended periods. These archives of preserved clinical FFPE specimens represent a significant, yet underutilized resource for advanced volumetric analysis.

Recently, the combination of high-resolution 3D imaging with advanced 3D bioimage analysis has emerged as a powerful toolkit in neuroscience [4, 16, 59, 60]. Thanks to the spread of automated image segmentation and classification methods, fluorescence imaging techniques, such as light-sheet fluorescence microscopy (LSFM) or two-photon fluorescence microscopy (TPFM), are evolving from purely observational to valuable quantitative analytical tools. Volumetric image reconstruction of human brain areas requires tissue optical transparency to reduce light scattering and absorption, thereby increasing the light penetration depth by refractive index matching [12, 46, 54, 56, 61]. Unlike for model organisms, clearing and labeling human brain tissues represents a challenging task. The major limitations are due to the variability in fixation and storage conditions of human samples, the high autofluorescence signal of blood vessels and lipofuscin, and the significant heterogeneity in cellular composition, structure, and optical properties across different regions and individuals [13, 28, 40, 43, 50, 58, 63]. To overcome some of these limitations, clearing methods such as CLARITY, SWITCH, SHIELD, iDISCO, and CUBIC, which were primarily developed for model organisms, have been adapted for human tissues [10, 38, 41, 45, 53, 55, 63]. Among the clearing methods specifically developed for large-scale human brain analysis, the SHORT tissue transformation method [14, 15, 43] enables rapid and efficient clearing of thick brain slices. SHORT results in high tissue transparency, structure preservation, reduction of intrinsic autofluorescence, and uniform sample co-staining by means of several cellular markers.

To harness the full potential of clearing techniques on human FFPE samples, an essential step in the workflow involves the full removal of the paraffin matrix from tissue blocks. Harsh deparaffinization methods are routinely used for proteomics or genomics analysis; however, these protocols require a homogenization step, either for proteins, DNA, or RNA extraction; which does not preserve the architecture of the sample [21, 32, 35, 42]. The development of volumetric reconstruction methods prompted researchers to optimize protocols of deparaffinization that maintain the structural organization of the sample, but the few alternatives proposed so far either for mouse or human tissues have been applied only to lungs, lymph nodes, and tumors [29, 39, 57]. The limited application of the deparaffinization step in combination with sample clearing and labeling methods highlights the difficulty in establishing a reliable protocol for paraffin removal from human brain tissues.

The ideal method for deparaffinization must strike a delicate balance, effectively removing the paraffin that could interfere with subsequent clearing steps, while preserving the integrity of tissue and protein structures. This is particularly difficult with human brain tissue since epitope preservation is essential for a homogeneous and specific labeling of the structures of interest, ensuring accurate visualization and analysis in downstream volumetric imaging.

Here we present a mild deparaffinization method that preserves the molecular and structural tissue architecture and works on blocks of human brain tissue of different sizes. This method was applied to adult postmortem human brain areas (brainstem), to postsurgical brain specimens removed from pediatric patients affected by malformations of cortical development (MCD) and to adult patients affected by hippocampal sclerosis (HS). Deparaffinized samples were processed with different clearing methods and reconstructed either with custom-made LSFM or TPFM setups.

Finally, to extract quantitative information on the spatial distribution and morphology of different classes of neurons from the 3D reconstructions of cleared samples, we implemented a machine - learning-based image segmentation workflow. This was employed to investigate the structural differences between two pathological samples of hemimegalencephaly (HME) and focal cortical dysplasia type IIa (FCDIIa). In detail, the cytoarchitectural organization of the samples was characterized by analyzing the neuronal volume, ellipticity, local density, and clustering level of two different neuronal populations.

Overall, we present a comprehensive approach for performing 3D imaging and automated quantitative cellular analysis on FFPE human brain specimens. This workflow includes human brain tissue deparaffinization, optical clearing, multi-labeling, LSFM or TPFM imaging, automated neuronal segmentation, and quantitative cytoarchitectural analysis (Fig. 1).

**Figure 1.**
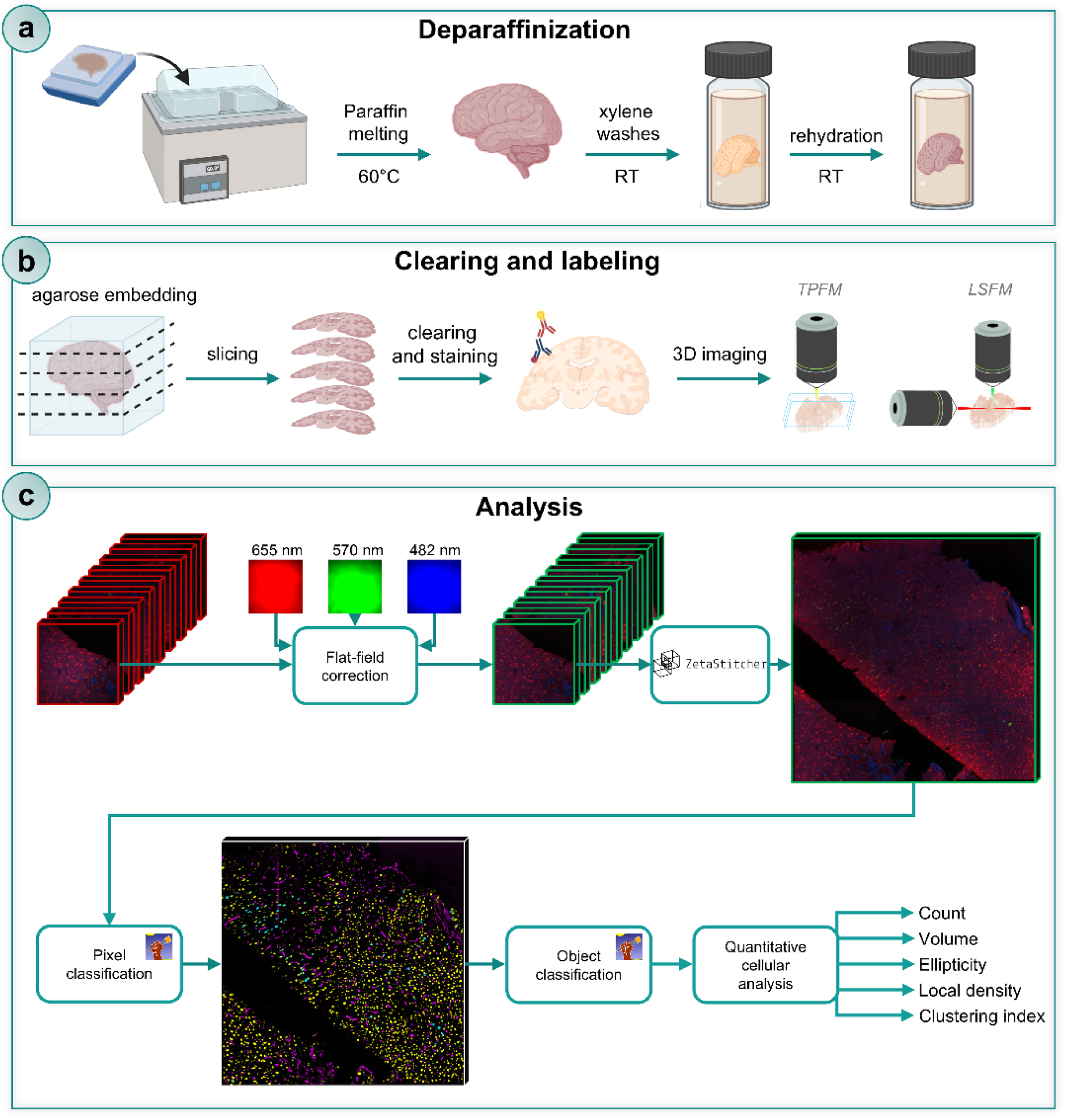
Overview of the entire pipeline for deparaffinization, clearing, labeling, imaging, automated neuronal segmentation and analysis. Schematic representation of the entire pipeline. a) Deparaffinization of FFPE (Formalin-Fixed Paraffin-Embedded) adult and pediatric human brain tissues. b) Clearing and labeling of deparaffinized slabs with the SHORT tissue transformation method, followed by volumetric imaging with LSFM (Light Sheet Fluorescence Microscopy) and TPFM (Two Photon Fluorescent Microscopy) custom-made setup. c) Block diagram of the 3D image processing pipeline for quantitative cytoarchitectural analysis of TPFM images. Adjacent overlapping TPFM image stacks first undergo flat-field correction using spatial gain models estimated with the CIDRE retrospective method. ZetaStitcher is then used to align the corrected stacks in order to create high-resolution image reconstructions of the brain specimens. Supervised pixel and object random forest classifiers, trained using the ilastik interactive machine learning tool, assess these reconstructions in sequence to automatically identify pRPS6+ and pRPS6-neuronal bodies. Finally, quantitative structural and morphological features are evaluated.

This approach grants access to the wide range of paraffin-embedded clinical specimens, enabling 3D quantitative analysis with cellular resolution and, thus, providing unprecedented insights into human brain cytoarchitecture in both physiological and pathological tissue samples.

## Results

### Deparaffinization and clearing of FFPE human brain tissue blocks

Formalin-fixed paraffin-embedded (FFPE) tissues are the most abundant type of samples available in neuropathology laboratories. To perform tissue clearing and 3D volumetric imaging, we optimized a method to completely remove the paraffin from human brain tissue blocks. We successfully applied our protocol to different types of brain samples from both healthy individuals and patients with malformations of cortical development (MCDs) and hippocampal sclerosis (HS), both pediatric and adult. In particular, we demonstrated the versatility of our method by processing post-mortem samples (brainstem) as well as samples surgically removed from an adult patient with HS and a pediatric patient with focal cortical dysplasia type IIa (FCDIIa) (Fig. S1). Deparaffinization was achieved by melting the paraffin wax surrounding the tissue block in a water-bath at 60°C followed by several washes in xylene, until the tissue block appeared completely translucent. These steps ensure the full removal of paraffin, both externally and deep inside the tissue. Following deparaffinization, xylene was removed with 100% ethanol, and the samples were rehydrated with a series of graded ethanol, until PBS was used for washes.

From the deparaffinized FFPE blocks, 500 μm-thick slices were cut and used to evaluate the compatibility of the deparaffinization protocol with two different clearing methods: the SHORT tissue transformation method [15, 43] and the iDISCO protocol [45]. To reconstruct the 3D structure of the cleared and stained volume, samples were imaged with a custom-made dual-view inverted LSFM setup [14, 33], with a isotropic final voxel size of 3.64 μm following post-processing.

In order to demonstrate the compatibility of the methodology with the SHORT technique and with multiple staining (e.g., targeting various neuronal subpopulations), we tested several markers (Table S1). The acquired LSFM volumetric reconstructions (Fig. 2) showed uniform and specific co-staining throughout the tissue depth. The slice from the brainstem was labeled with two markers to characterize different subpopulations of GABAergic interneurons: Somatostatin (SST) and Calretinin (CR) (Fig 2a, 2b, 2c). Whereas the slab from the surgical hippocampus (HS-affected patient) was labeled with the NeuN antibody along with the nuclear marker DAPI. The 3D reconstructions obtained with LSFM revealed a specific staining of neurons across various hippocampal regions (Fig. 2d, 2e, 2f). As expected, we observed segmental pyramidal cell loss in several Cornu Ammonis (CA) sectors (Fig. S2a, S2b), typical histopathologic hallmark of HS [6], as confirmed in FFPE tissue sections from the same block. Immunohistochemistry and histological staining on adjacent paraffin blocks confirmed this finding (Fig. S2c, S2d, S2e and S2f). Finally, the FCD brain slab was marked with the pan-neuronal marker NeuN and the phospho-ribosomal protein S6 (pRPS6^Ser235/236^) (Fig. 2g, 2h, 2i). The latter was used as a readout of the phosphatidylinositol 3-kinase (PI3K)/AKT/mammalian target of the rapamycin (mTOR) pathway dysregulation, which characterizes the disease [34].

**Figure 2.**
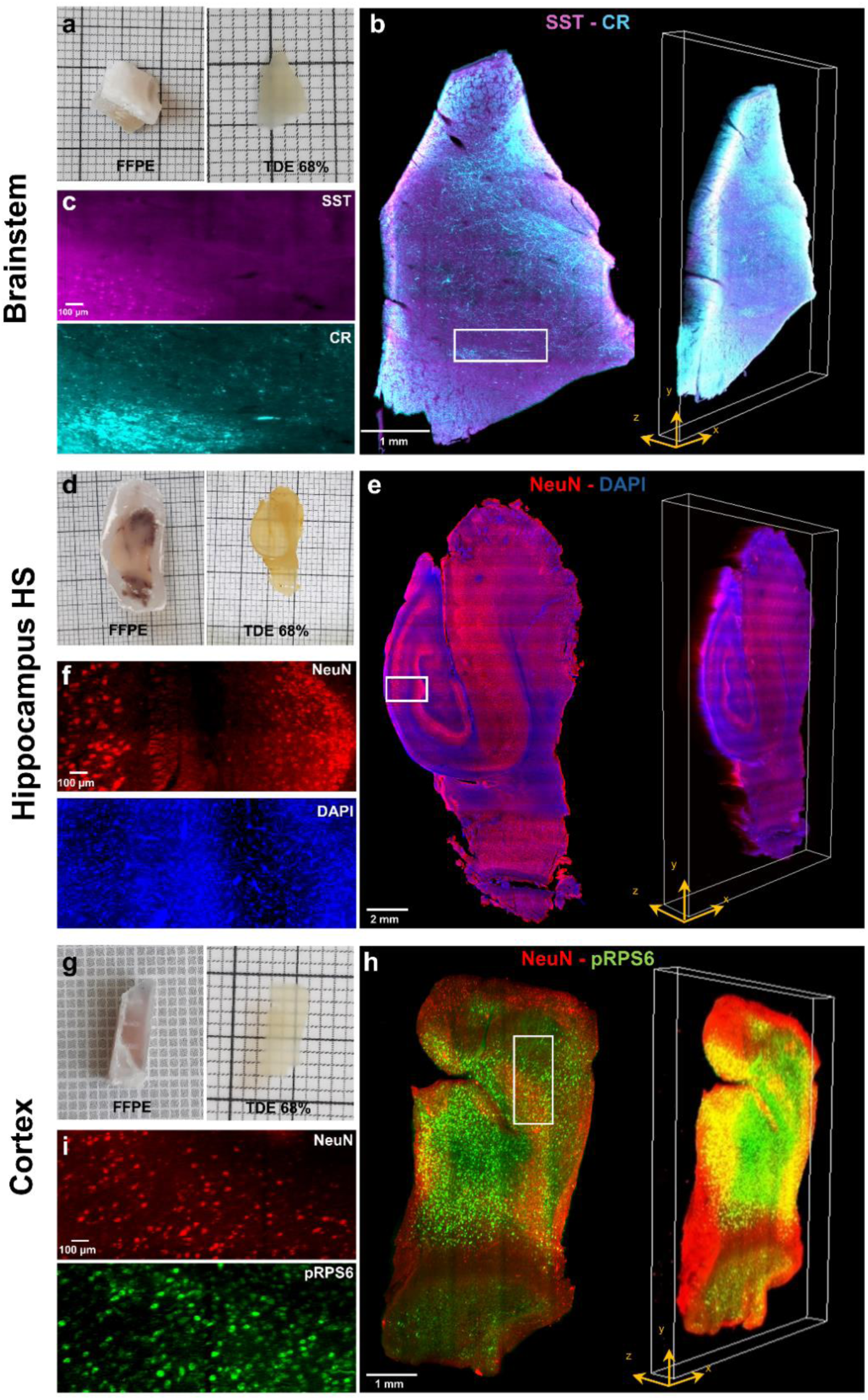
3D reconstructions with LSFM of deparaffinized human brain slabs processed with the SHORT tissue transformation method. **a)** Image of a postmortem adult human brainstem, before deparaffinization (FFPE) and after SHORT (TDE 68%)**. b)** Maximum intensity projection image (3.64 µm isotropic resolution) and volumetric rendering showing a mesoscopic reconstruction of (a) labeled for Somatostatin (SST, magenta) and Calretinin (CR, cyan). Scale bar: 1 mm. **c)** Insets of (b) showing the single markers used, SST, (magenta) and CR (cyan). Scale bar: 100 µm. **d)** Image of a postsurgical human hippocampus from a patient with hippocampal sclerosis (HS), before deparaffinization (FFPE) and after SHORT (TDE 68%)**. e)** Maximum intensity projection image and volumetric rendering showing a mesoscopic reconstruction of (d) labeled for NeuN (red) and DAPI (blue). Scale bar: 2 mm. **f)** Insets of (e) showing the single markers used, NeuN (red) and DAPI (blue). Scale bar: 100 µm. **g)** Image of a postsurgical brain specimen from a pediatric patient with focal cortical dysplasia type IIa (FCDIIa), before deparaffinization (FFPE) and after SHORT (TDE 68%)**. h)** Maximum intensity projection image and volumetric rendering showing a mesoscopic reconstruction of (g) labeled for NeuN (red) and pRPS6 (green). Scale bar: 1 mm. **i)** Insets of (h) showing the single markers used, NeuN (red) and pRPS6 (green). Scale bar: 100 µm. The insets in (c), (f), and (i) refer to the regions in white boxes in (b), (e) and (h).

To assess the compatibility of the deparaffinization protocol with different clearing methods, we also employed it in combination with the organic solvent-based technique iDISCO [45]. Using this technique, we achieved high levels of tissue transparency and multi-staining with selected markers (Table S1) in different human brain specimens: a postmortem adult brainstem (Fig. S3a, S3b), a postsurgical hippocampus (HS-affected patient) (Fig. S3c, S3d), and a pediatric FCD-affected cortical brain sample (Fig. S3e, S3f). However, high levels of natural autofluorescence from the lipofuscin-type pigments were visible in the adult samples when imaging with excitation wavelengths with two distinct clearing methods: the SHORT tissue transformation technique and the organic solvent-based iDISCO. This entire process was applied to adult and pediatric specimens from various brain regions, enabling 3D reconstruction of the samples acquired with LSFM at micrometer resolution.

### Quantitative cytoarchitectural analysis on deparaffinized pathological human brain samples

The successful deparaffinization, clearing, and co-labeling of pathological human brain samples prompted us to evaluate and compare quantitative cytoarchitectural features, namely local neuronal density and clustering - a structural parameter shown to have a marked functional connotation [17, 18] - along with morphological cell properties, such as soma volume and ellipticity. To this aim, mesoscopic image reconstructions of 500-μm-thick sections with a resolution of 1.21 µm × 1.21 µm × 2 µm were acquired using a two-photon fluorescence microscope (TPFM) (Fig. 3). Two brain specimens surgically removed from pediatric patients were examined: a sample from a patient affected by HME, (S_1_) and one by FCDIIa, (S_2_) (Fig. 3a, 3b). These two patients underwent genetic testing. In patient S_1_, massive parallel sequencing of a panel containing 48 genes of the mTOR pathway revealed the heterozygous c.392C>T (p.Thr131Ile) variant of uncertain significance (VUS) in the *PTEN* gene (NM_000314.4), which was inherited from the healthy mother. Patient S_2_ corresponds to Patient 1 described by Guerrini and collaborators [23] and carried the c.6644C>T (p.Ser2215Phe) MTOR (NM_004958.3) pathogenic variant, which was present at 5.5% alternative allele fraction (AAF) (GS Junior sequencing validation: 2.46%) in the dysplastic brain tissue and absent in DNA extracted from saliva and blood.

**Figure 3.**
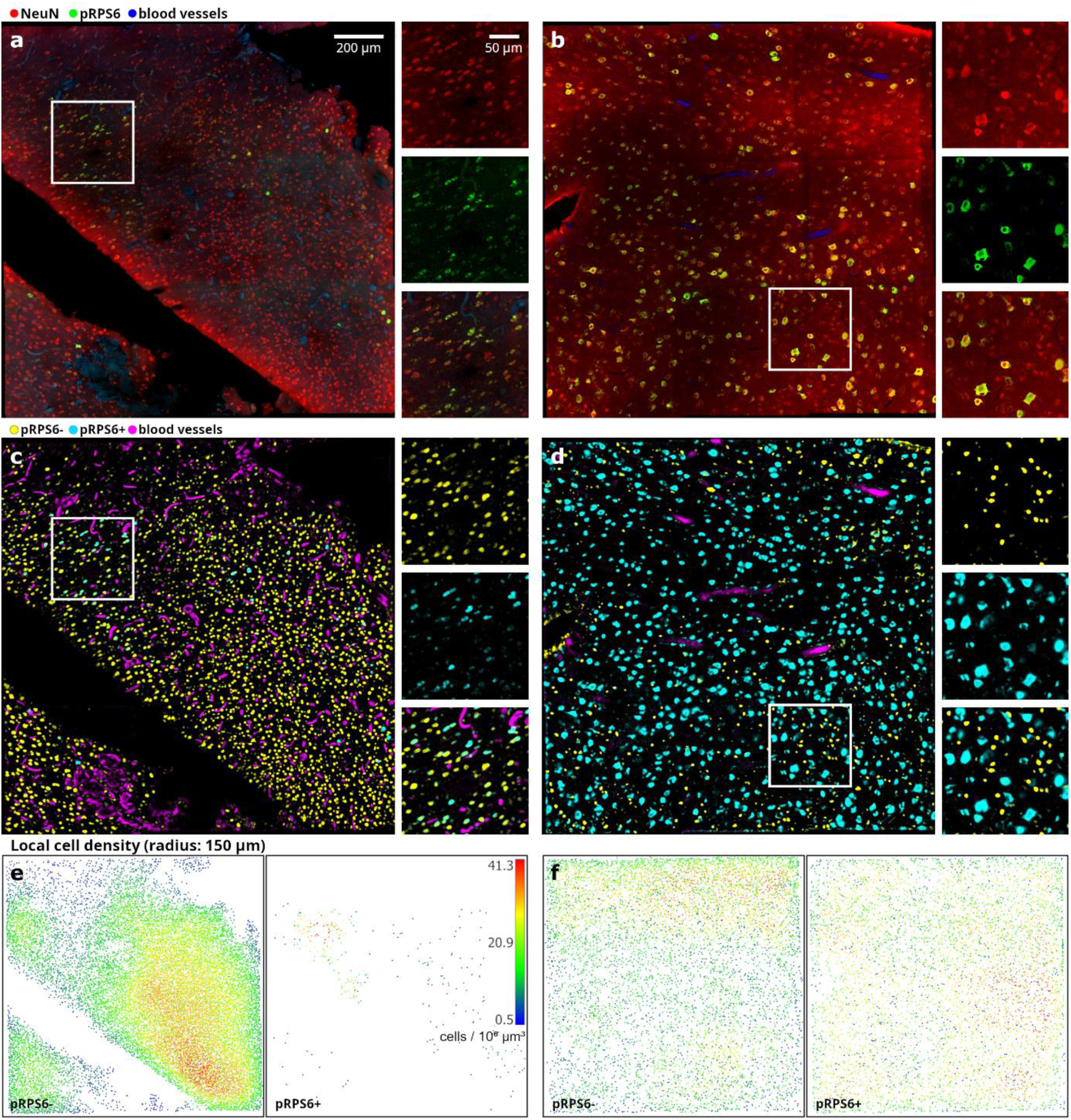
Volumetric imaging with TPFM and analysis on surgically removed pediatric brain specimens. **a-b)** Representative middle plane of mesoscopic reconstructions obtained with TPFM (1.2 μm × 1.2 μm × 2 μm resolution) are shown for surgically removed brain pieces from patients affected by hemimegalencephaly (S_1_, a) and focal cortical dysplasia type IIa (S_2_, b). Tissues were labeled for NeuN (red) and pRPS6 (green) while blood vessels were detected through autofluorescence (blue). Scale bar: 200 µm. On the right the magnified insets showing the single markers used. Scale bar: 50 µm. The insets images refer to the regions in white boxes. **c-d)** Corresponding 3D semantic segmentations generated using ilastik’s pixel classification workflow (headless prediction on a distributed computer cluster). Yellow: pRPS6-neurons; cyan: pRPS6+ neurons; magenta: blood vessels. Background suppression shows striking accuracy. **e-f)** Local cell density maps respectively generated from the TPFM reconstructions in a-b (left: pRPS6-neurons; right: pRPS6+ neurons).

Specific cytoarchitectural alterations in the population of neurons expressing the phospho-Ribosomal Protein S6^Ser235/236^ (pRPS6+) were quantitatively assessed with respect to neurons stained with anti-NeuN antibody through machine-learning-based image enhancement and automatic cell segmentation (Fig. 3c, 3d). In particular, semantic segmentation of TPFM image reconstructions, automatic recognition and quantitative characterization of neuronal bodies were carried out using ilastik [4], a user-friendly, open-source software for bioimage analysis. Leveraging the power of supervised machine learning, ilastik simplifies the often complex and labor-intensive task of image segmentation by allowing users to interactively train random forest image classifiers (Breiman L., 2001) through a highly intuitive interface. Despite being considerably slower than state-of-the-art deep learning-based tools for automatic cell centroid localization, such as the GPU-implemented BCFind-v2 [8], ilastik enables the faithful semantic segmentation of neuronal bodies and, thus, the quantitative evaluation of their volume and shape. In this work, automatic neuron detection was achieved by means of two separate machine learning workflows supported by ilastik, i.e. pixel and object classification. Following the interactive manual training, both ilastik’s workflows were run in headless mode on a high-performance cluster comprising four computing nodes. An average out-of-bag error [26] of 2.8% was achieved on the training set generated by the three expert operators involved in the manual image annotation task. This was mirrored in the precise automatic detection of both pRPS6+ and pRPS6-cell bodies in their respective probability channels, as well as in the faithful 3D reconstruction of local microvascular trees, thanks to accurate tissue background suppression. The subsequent object classification stage further improved of the raw detections, enabling us to robustly distinguish the presence of multiple tightly packed cells that had been unavoidably merged during the binarization of the semantic maps returned by the upstream image enhancement workflow. This was crucial for preventing local underestimates of the actual cellular density and thus for the correct representation of spatial and inter-subject variations in the brain tissue cytoarchitecture. To visualize the cluster of pRPS6+ neurons in the HME tissue specimen (S_1_) and the remarkably higher prevalence of pRPS6-expressing cells observed in FCDIIa (S_2_) we obtained neuronal density maps for both the subjects (Fig. 3e, 3f).

The method detected pRPS6-expressing neurons in both tissue samples but the percentage of pRPS6+ neurons relative to the total number of neurons, identified by the NeuN staining, resulted remarkably higher in S_2_ (35.4%) compared to S_1_ (3.1%) (Table S2). This difference is in line with the finding that genetic testing revealed only a probably benign VUS in S_1_, while S_2_ was found to carry a mTOR pathogenic variant that has been reported to cause strong hyperactivation of the mTOR pathway [23].

Interestingly, in S_2_ we also observed a statistically significant increase in the neuronal volume (Bonferroni-corrected p < 0.001; Hellinger distance, H = 0.308, inter-median distance, |ΔM| = 2884.3 μm^3^; Fig. 4a and 4b) and ellipticity (p < 0.001, H = 0.282, ΔM = 0.4; Fig. 4c and 4d) of pRPS6+ neurons, compared to S_1_. This morphological alteration pattern resembles the one already highlighted by neuropathological evaluations of FCD brain tissue, that revealed the presence of dysmorphic giant neurons exhibiting RPS6 hyperphosphorylation [23]. S_2_ also showed a slight increase in the cell body volume and ellipticity of neurons not expressing the pRPS6 protein (pRPS6-), compared to S_1_ (p < 0.001, H = 0.154, |ΔM| = 579.7 μm^3^; p < 0.001, H = 0.122, |ΔM| = 0.1; Fig. 4a-d). This finding may be ascribed to the differences in cell types and distribution in different brain areas, as tissues used in this study have been sampled from the parietal lobe for S_1_ and the frontal lobe for S_2_. We also detected statistically significant differences between S_1_ and S_2_ regarding their local neuronal density (within a radius of 150 μm). In detail, compared to S_1_, S_2_ exhibited a lower cell density of neurons not expressing pRPS6 (p < 0.001, H = 0.606, |ΔM| = 11 cell / 10^6^ µm^3^; Fig. 4e and 4f) and a higher density of pRPS6+ cells (p < 0.001, H = 0.612, |ΔM| = 4 cell / 10^6^ µm^3^; Fig. 4e and 4f). Thus, the pronounced increase in the percentage of pRPS6-expressing neurons reported above for subject S_2_. Differences in cell distribution may also explain the modest but statistically significant difference between S_1_ and S_2_ that emerged for the clustering indices of pRPS6-neurons (p < 0.001, H = 0.192, |ΔM| = 0.05 a.u.; Fig. 4g and 4h). On the other hand, it is more relevant to emphasize the peculiar distribution of the clustering indices of pRPS6+ neurons in subject S_1_ compared to that of S_2_ (Fig. 4g and 4h). Indeed, differently from the narrow data distributions observed in S_2_ (indicating, to a greater or lesser extent, a uniform spatial arrangement across the acquired TPFM image reconstructions), in S_1_ the pRPS6+ cell clustering indices show a wide distribution spreading from negative data with a marked peak at the lowest value of the distribution, specifically associated with the presence of completely isolated cells, to high values corresponding to local clusters of neurons. This manifests in statistically significant differences with respect to the pRPS6+ clustering data distribution of subject S_2_ (p < 0.01, H = 0.332, |ΔM| = 0.05 a.u.; Fig. 4h) and to the reference clustering indices of neurons pRPS6-evaluated for that subject (p < 0.01, H = 0.453, |ΔM| = 0.07 a.u.; Fig. 4h). All statistical results are reported in Table S3.

**Figure 4.**
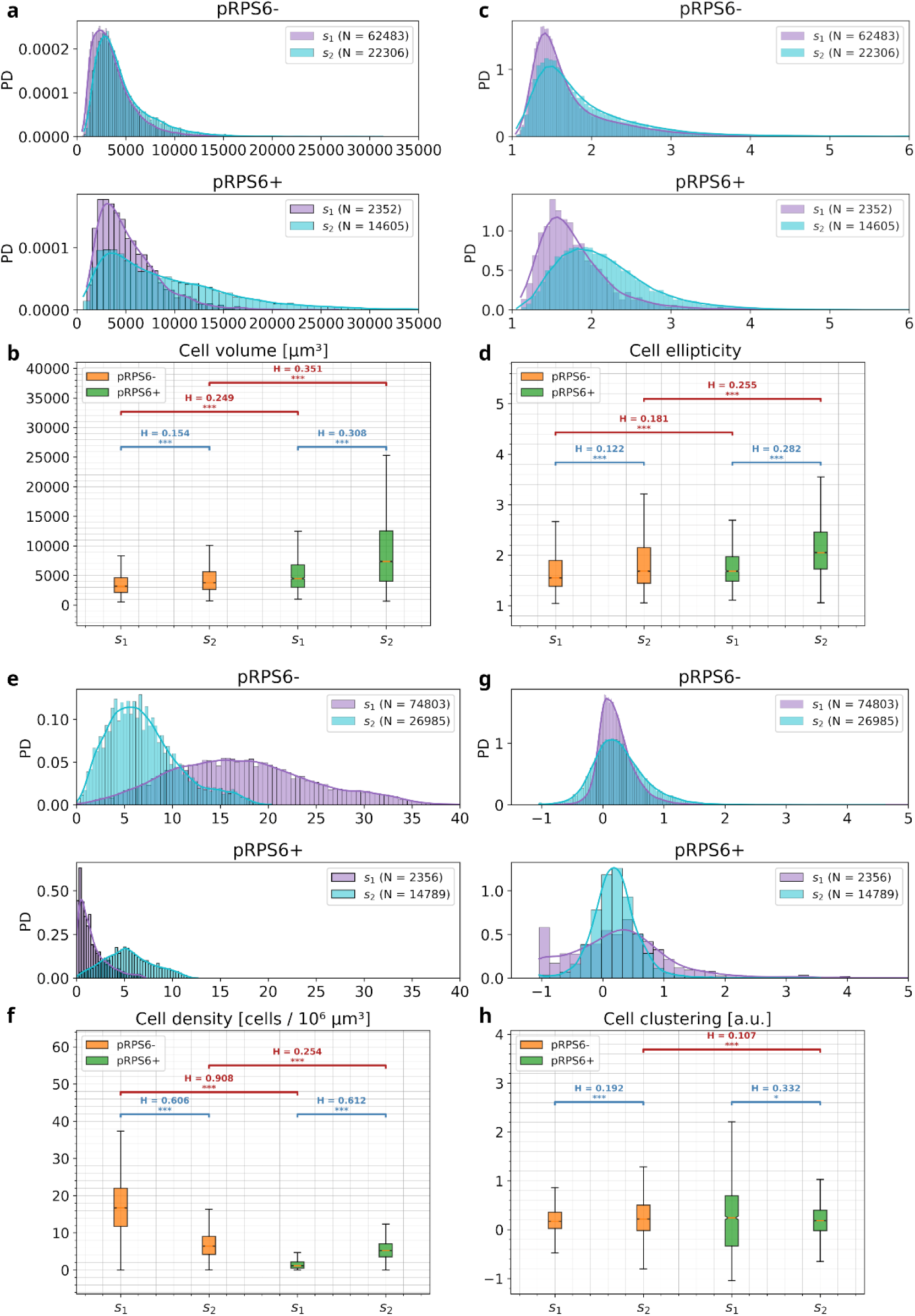
Quantitative observed cytoarchitectural analysis on pediatric human brain specimens. Quantitative cytoarchitectural analysis of neuronal body morphology (a, b: cell volume; c, d: cell ellipticity) and spatial organization (e, f: local cell density; g, h: clustering index). PD: probability density; *** p < 0.00017, ** p < 0.00167, Mann -Whitney U-Test (Bonferroni correction); H: Hellinger distance between the compared data distributions.

## Discussion and conclusions

Volumetric reconstruction with sub-cellular resolution can provide detailed insights into the complexity of human brain architecture and pathology. In this work, we introduce a comprehensive workflow to perform 3D reconstruction and automated quantitative cellular analysis on deparaffinized human brain samples. We developed a versatile deparaffinization protocol for human brain tissue blocks, compatible with different optical clearing and multi-labeling methods. Volumetric reconstructions were analyzed to extract quantitative information about the morphology and spatial distribution of different classes of neurons, through a machine-learning-based image segmentation workflow (Fig. 1).

Optical clearing methods are widely used on fresh or archival formalin-fixed tissues. However, a substantial fraction of human samples stored in biobanks are embedded in a paraffin matrix to preserve tissue integrity. Efficient clearing and 3D imaging require paraffin removal, which otherwise acts as a physical barrier, impeding the diffusion of clearing agents, thus leading to uneven optical transparency that compromises the image quality. Residual paraffin can also hinder the accurate labeling of structures of interest. Harsh deparaffinization methods, such as prolonged exposure to organic solvents or high temperatures, can induce epitope masking effects and/or antigen damage, affecting the accuracy of subsequent analyses.

The mild deparaffinization method proposed here allows for the complete removal of paraffin from human brain tissue blocks by combining a water bath’s wet heat to eliminate outer paraffin and multiple xylene incubations for inner paraffin removal. Unlike the approaches presented for murine models, for human brain tissue it was necessary to increase the frequency and reduce the duration of the single xylene washes, performed using a rotatory shaker. This approach ensured the complete removal of inner paraffin while preserving the tissue’s macrostructure and antigenicity, as demonstrated by subsequent stainings.

There are significant differences in tissue architecture and optical properties between adult and pediatric brain samples, as well as between tissues from various regions of the human brain [1, 2, 19, 27, 36]. Additionally, factors such as the duration of formalin fixation, and pre- and postmortem changes, such as pH variation and tissue oxygenation, vary significantly from one brain to another. Reflecting its versatility, our deparaffinization protocol effectively enabled the removal of paraffin from different specimens, including normal and abnormal patient-derived human brain samples from both adult and pediatric cases. Two methodologies were tested for clearing and labeling: the SHORT tissue transformation technique [15, 43] and the organic solvent-based iDISCO [45]. Both methods achieved high tissue transparency and specific, homogeneous co-staining with multiple antibodies, allowing us to visualize different neuronal populations and characterize the pathological samples. However, the SHORT technique was more effective at quenching lipofuscin autofluorescence and proved more efficient for labeling pediatric samples with respect to iDISCO. Therefore, the quantitative evaluation on neuronal subpopulations was performed on SHORT-cleared slabs.

Neuronal and glial cell numbers are important attributes in the characterization of distinct functional regions of the nervous system. These numbers are moreover susceptible to pathologies, pharmacological treatments, and genetic alterations [20, 25, 30, 44]. The vast majority of reports on cell counts in the neuroscience literature are based on stereological methods [22, 49, 62]. In these approaches, the number of cells is accurately measured in a small but unbiased portion of the volume of interest and the numbers for the whole target region can then be extrapolated under the assumption that the sample is representative. Since stereological neuron counting requires the involvement of a human operator to identify cells in the stained tissue [48], the procedure is inherently laborious, time-consuming, and in particular for cell-sparse regions, rather inefficient [7].

However, in recent years, image analysis has been revolutionized by machine learning (ML). Thanks to the growing adoption of ML-based image segmentation, 3D fluorescence imaging techniques, such as light-sheet fluorescence microscopy (LSFM) and two-photon fluorescence microscopy (TPFM), are nowadays evolving from being just a qualitative observational modality to a valuable analytical tool that enables the automatic extraction of quantitative cytoarchitectural information from 3D mesoscopic reconstructions of tissue samples imaged with sub-cellular resolution. This was so far unfeasible to perform for human operators with adequate reliability and within practical time frames. The manual labeling effort required to robustly train a supervised image classification algorithm with sufficiently rich image feature data may, however, still be considerable, if not impossible. In this regard, the adoption of the ilastik toolkit [4] for interactive ML-based image analysis has proved of the utmost importance, providing us with a convenient user interface supporting fast interactive training with immediate feedback on the achieved segmentation accuracy. Indeed, trained pixel and cell body classifiers were ready within a few working days. This capability to swiftly deploy and execute ML-based bioimage analysis workflows, as shown in the present work, will be crucial to building meaningful large-scale human brain cytoarchitectural datasets for future neuroanatomical studies.

The superior resolution of the TPFM setup prompted the decision to perform the quantitative evaluations on mesoscopic reconstructions obtained with this system from two surgically removed pediatric brain specimens, previously imaged by LSFM. Since genetic testing on these specimens revealed mutations in genes belonging to the mTOR pathway (PTEN for the HME tissue and MTOR for the FCDIIa tissue), we selected the cell populations to analyze based on the expression of the pRPS6, a marker of mTOR signaling activation, overexpressed in most MCDs [34]. We first evaluated the percentage of pRPS6+ neurons relative to the total number of neurons identified by the NeuN staining and found it to be significantly higher in the FCDIIa (S_2_) sample (∼35%) than in the HME (S_1_) sample (∼3%). In line with this finding, Patient S_2_ carries a pathogenic variant in the MTOR gene, previously reported to cause a strong hyperactivation of the mTOR pathway [23]. The low percentage of pRPS6+ neurons observed in the HME tissue could be ascribed to the fact that the VUS in PTEN identified in Patient S_1_ might be benign, with a slight effect on mTOR pathway activation. The downstream signaling of PTEN can be mediated by two complexes mTORC1 and mTORC2, which have different downstream targets. Considering that RPS6 is a downstream effector of the mTORC1 complex, we cannot rule out that the PTEN variant in Patient S_1_ may be pathogenic and exerts its effect independently of pRPS6, by preferentially activating the mTORC2 complex. Selective deregulation of mTORC2 caused by mutant PTEN has been previously documented in a model of human glioblastoma cell lines [5] and in PTEN-deficient mice [9]. Our analysis can detect also morphological alterations: a statistically significant increase in the neuronal volume and ellipticity of pRPS6+ neurons in FCDIIa compared to HME was found, highlighting a well-known characteristic of FCD brain tissue that exhibits a high presence of dysmorphic giant neurons were RPS6 is hyperphosphorylated. Additionally, we observed interesting differences in cell clustering indices: in HME, pRPS6+ cells displayed a broad distribution, including both isolated neurons and local clusters, whereas in FCDIIa, pRPS6+ cells exhibited a uniform, non-clustering distribution. Future studies are needed to further investigated the possible correlation of this pRPS6+ cells clusters with epileptogenic foci, as well as the variance of cell density distributions in MCDs. Nevertheless, the protocol we set up provide new insights in clarifying pathophysiological alterations of brain anomalies, such as focal MCDs. In fact, most data used to define histopathological alterations of FCD and HME come from bidimensional (2D) images, but such images do not allow for the reconstruction all brain networks altered in these conditions. Cells carrying pathogenic variants may establish connections with others - either mutant or WT - located on different focal planes. Thus, tracing the connections of mutant cells along different focal planes will improve our knowledge of how they interact with the surrounding tissue. The protocol designed has also the potential to disclose new morphological cell markers that may impact the diagnosis and classification of focal MCDs.

In conclusion, our workflow offers valuable tools for investigating human brain tissues, highlighting the untapped potential of FFPE tissue archives for advanced volumetric analysis. This could help in clarifying pathophysiological alterations and uncover new morphological markers, potentially impacting the diagnosis and classification of neurological diseases.

## Materials and methods

### Samples

The study was approved by the Pediatric Ethics Committee of the Tuscany Region (Italy) in the context of the DESIRE FP7 EU project and its extension by the DECODE-EE and Human Brain Optical Mapping projects. Pediatric human brain samples were removed during surgical procedures for the treatment of drug-resistant epilepsy in children with focal MCDs. Samples were obtained after informed consent, according to the guidelines of the Pediatric Research Ethics Committee of the Tuscany Region. Upon collection, samples were placed in neutral buffered formalin (pH 7.2–7.4) (Diapath, Martinengo, Italy) and stored at 4°C until the deparaffinization and clearing process.

Healthy human brainstem tissue was obtained from the Body Donation Program “Donation to Science” at the University of Padova. Before death, participants provided their written consent for the use of their entire body for any educational or research purpose in which the anatomy laboratory is involved. The authorization documents (in the form of handwritten testaments) are kept in the files of the Body Donation Program. Upon collection, samples were fixed in 4% formalin solution, embedded in paraffin and stored at room temperature (RT) until the deparaffinization and clearing process.

The human hippocampus with histopathological features of HS was obtained from an adult patient who underwent surgery as treatment for drug-resistant epilepsy at the Fondazione Istituto Neurologico Carlo Besta (Milan, Italy). The resection was performed for strictly therapeutic reasons, after informed consent, and the extent of the excision was planned preoperatively based on both the epileptogenic zone localization and the risk of postsurgical deficits. The local ethics committee approved the use of brain material for research purposes.

### Deparaffinization of brain tissue

Deparaffinization was achieved by first removing as much paraffin surrounding the tissue block as possible with a blade. To completely remove the remaining paraffin, samples were placed in a water bath at 60°C for 30-60 min until the paraffin was macroscopically eliminated. In order to speed up this step and completely clean samples from the paraffin, tissue blocks were positioned on a porous membrane inside a 50 ml tube floating in the water bath. Following the paraffin melting, tissue blocks were incubated 6-8 times in 98% Xylene for 30 minutes at RT, using a rotary shaker. At this point, tissue blocks should appear translucent. Afterwards, samples were washed with 100%, 95%, and 70% ethanol diluted with water at RT for 30 min respectively, and further washed 3 times with 1× PBS for 10 min at RT. After embedding in 4% agarose, tissue sections of 500 ± 50 μm-thick were obtained using a vibratome (Leica VT1000 S) and stored at 4°C in PBS + 0.01% w/v NaN_3_.

### SHORT tissue transformation method

Single deparaffinized slices with a thickness of 500 μm were processed with the SHORT tissue transformation method, as previously described [15, 43]. The specimens were first incubated in a Switch-Off solution (50% v/v 1× PBS (pH 3), 25% v/v 0.1 M hydrochloric acid (HCl), 25% v/v 0.1 M potassium hydrogen phthalate (KHP), and fresh 4% v/v glutaraldehyde) at 4°C in gentle shaking for 24 h. The solution was replaced with the Switch-On (1× PBS (pH 7.4) and fresh 1% v/v glutaraldehyde) and the incubation was performed at 4°C in gentle shaking for 24h. Afterwards, slices were washed 3 times for 2 h each in 1× PBS at RT and reactive glutaraldehyde was inactivated by o/n (overnight) incubation with an inactivation solution (1× PBS (pH 7.4), 4% w/v acetamide, 4% PBS at RT and then incubated in the clearing solution (200 mM sodium dodecyl sulfate (SDS), 20 mM sodium sulfite (Na_2_SO_3_), and 20 mM boric acid (H_3_BO_3_), pH 9.0) for 3-6 days (Table S4) at 55°C. After the clearing step, samples were extensively washed in 1× PBS + 0.1% Triton X-100 (PBST) at 37°C for 24 h. Next, the slices were treated with 30% H_2_O_2_ for 45 min and antigens were unmasked with the preheated antigen retrieval solution (10 mM Tris base (v/v), 1 mM EDTA solution (w/v), 0.05% Tween 20 (v/v), pH 9) for 10 min at 95°C. After cooling down to RT, the specimens were washed in deionized (DI) water for 5 min each and then equilibrated with PBS for 1 h. The primary antibodies were diluted in PBST + 0.01% w/v NaN_3_ (Table S1 and S4), and sample incubation was performed at 37°C in gentle shaking. After washing with PBST at 37°C for 24 h, the stained samples were incubated in gentle shaking with the secondary antibodies at 37°C in PBST + 0.01% w/v NaN_3_ (Table S1 and S4). Next, the stained samples were extensively washed with PBST at 37°C, equilibrated first in 30% TDE/PBS (v/v), then in 68% TDE/PBS (v/v), and finally placed in the sandwich with 68% TDE.

### iDISCO method

Single deparaffinized slices with a thickness of 500 μm were processed with a modified iDISCO protocol [45]. Specimens were dehydrated with increasing concentrations of methanol MeOH/H_2_O (20%, 40%, 60%, 80%, and 100%), with gentle rotation for 1 h each at RT. Samples were then incubated in 66% dichloromethane (DCM) / 33% MeOH overnight at RT with gentle rotation. Afterwards, samples were washed with 100% MeOH for 1 h twice, and autofluorescence was bleached through incubation with 5% H_2_O_2_ in MeOH at 4°C overnight, without shaking. After bleaching, samples were rehydrated with decreasing concentrations of MeOH/H_2_O (80%, 60%, 40%, 20%), followed by one PBS and two PBST (1× PBS and 0.2% v/v Triton X-100) washes, all with gentle rotation for 1 h each at RT. On the same day, samples were incubated first with the Permeabilization solution (1× PBS, 0.2% v/v Triton X-100, 20% v/v DMSO and 0.3M glycine), followed by incubation in Blocking solution (1× PBS, 0.2% v/v Triton X-100, 0.2% w/v gelatin porcin skin and 0.01% w/v NaN_3_), both for 24 h, at 37°C with gentle rotation. Primary antibodies (Table S1) were diluted in PBSTW-Heparin (1× PBS, 0.2% v/v TWEEN-20 and 0.01 mg/ml Heparin) for 6 days, at 37°C with gentle rotation. Samples were then washed for 1 day in PBSTw-Heparin and then incubated with the secondary antibodies (Table S1) in PBSTw-Heparin for 4 days, at 37°C with gentle rotation. After 1 day of washes in PBSTw-Heparin, samples were dehydrated as previously described and incubated for 3 h in 66% DCM / 33% MeOH. Finally, they were washed in 100% DCM for 30 min twice and incubated and stored in DBE (Dibenzyl ether) at RT.

### TPFM imaging

A custom-made TPFM system [11, 52] was used to acquire high-resolution 3D image reconstructions of the brain tissue slices. The system employs a Chameleon (Coherent, US) tunable mode-locked Ti:Sapphire laser (pulse rate: 90 MHz, pulse width: 120 fs) as the excitation source. The light beam is laterally scanned by means of an optically coupled pair of galvanometric mirrors (LSKGG4/M, Thorlabs, US) and focused onto tissue samples using a 20× objective (Nikon CFI90 20XC Glyc, JP; numerical aperture (NA): 1.0, working distance (WD): 8.2 mm), with a resulting field of view of 621 μm × 621 μm. Two-photon excitation was achieved by operating the laser source in the near-infrared region at a wavelength of 800 nm. 655/40 nm, 618/50 nm, and 482/35 nm single-band bandpass filters (BrightLine®, Semrock, US) were respectively used to detect the fluorescence emission of neuronal bodies (NeuN), pRPS6+ neurons, and blood vessels (second harmonic autofluorescence). The filtered fluorescence emission is collected and amplified by three GaAsP photomultiplier tubes (H7422, Hamamatsu Photonics, JP), and digitized with 8-bit precision. The sequential imaging of adjacent overlapping 3D stacks is enabled by a closed-loop XY stage (U-780 PILine XY Stage System, Physik Instrumente, GER) for the lateral translation of tissue specimens, and a closed-loop piezoelectric stage (ND72Z2LAQ PIFOC Objective Scanning System, Physik Instrumente, GER) for the axial displacement of the objective. This, in turn, allows for the acquisition of high-resolution tiled image reconstructions of entire brain slices. The nominal lateral overlap of approximately 50 μm between adjacent tiles, adopted in the present work, ensured an optimal fusion with a precise superimposition of corresponding tissue structures. In this study, 512 × 512 pixel images were acquired with a z-step of 2 μm, resulting in a final voxel size of 1.21 μm × 1.21 μm × 2 μm. The uneven illumination (vignetting) of the separate images was corrected via the retrospective CIDRE method (Smith et al., 2015), in order to prevent the grid-like shading artifacts that would otherwise arise when fusing the adjacent stacks. The flat-field intensity models related to the wavelengths of interest (655 nm, 570 nm, and 482 nm) were identified using a reference dataset comprising 7630 uncorrelated RGB images acquired with the employed objective.

### LSFM imaging

A custom-made inverted light-sheet fluorescence microscopy (LSFM) setup described in a previous work of our group [14, 33] was also used to image the brain tissue samples. This custom LSFM setup includes two identical 12× objectives (LaVision Biotec LVMI-Fluor 12× PLAN, GER; NA: 0.53, WD: 8.5 to 11 mm), oriented at 90° relative to each other and spaced so that their FOVs overlap in the center. These objectives are inclined at 45° with respect to the horizontal sample holder plane and alternately play excitation and detection roles. The microscope is equipped with four laser diode modules (Cobolt, SE) characterized by the following wavelengths and nominal output powers: (405 nm, 100 mW); (488 nm, 60 mW); (561 nm, 100 mW); and (638 nm, 180 mW). These excitation sources emit Gaussian beams which are adjusted in width by dedicated telescopes, before being combined by means of three dichroic mirrors. This combined beam is then divided by a 50:50 beam splitter. Outgoing light is then introduced in two identical excitation pathways in which each beam is firstly modulated in intensity, phase, and wavelength using an acousto-optical tunable filter (AOTF, shuttering of each illumination pathway to avoid introducing stray light originating from the excitation beam during image acquisition. Secondly, the beam is scanned by means of a galvo mirror (6220H, Cambridge Technology, US) and then conveyed into a scan lens (#45-353, Edmund Optics, US; focal length (FL) = 100 mm, achromat) producing the planar light-sheet illumination. The generated light sheet is finally directed to the pupil of the 12× objective through a tube lens (#45-179, Edmund Optics, US; FL = 200 mm, achromat), to illuminate a single plane of the tissue sample. Each objective is mounted on a motorized stage (PI L-509.14AD00, Physik Instrumente, GER) in order to move its focal plane position. On the other hand, the sample is translated using a 3-axis motorized stage system (two PI M-531.DDG and a PI L-310.2ASD, Physik Instrumente, GER; overall motion range: 30 cm × 30 cm × 2.5 cm) and imaged while being shifted along the horizontal x-axis, through the fixed FOVs of the two objectives. The fluorescence emitted by the sample and collected by the detection objective is separated from the reflected excitation light by means of a multiband dichroic beamsplitter (Di03-R405/488/561/635-t3-55×75, Semrock, US) before being directed by the detection tube lens onto an sCMOS camera (Orca-Flash4.0 v3 Hamamatsu, JP; pixel size: 6.5 μm × 6.5 μm, pixel array: 2048 × 2048). Five sets of band-pass filters were mounted in front of each camera on a motorized filter wheel (FW102C, Thorlabs, US) to selectively image the differently labeled structures within tissue samples. Cameras operate in confocal detection mode by having the rolling shutter sweep in synchrony with the galvo scan of the light sheet [37, 51]. To prevent excitation stray light from interfering with the active rolling-shutter rows on the acquiring camera, a time delay of half a frame is introduced between the detection and excitation modes of each optical path. Following the completion of an image stack, the tissue sample is shifted along the y-axis (by a range below the FOV side: ≈1109 μm), and the sequence is repeated until the whole tissue volume has been acquired. In this work, a y-step of 1 mm was adopted for a lateral image overlap of more than 100 μm. An x-step of 5.15 μm, instead, was set to obtain a native voxel depth of 3.64 μm.

### Microscopy image post-processing

Since the cameras of the custom-made LSFM setup acquire images of brain tissue samples while these pass through the focal plane of the detection objectives with a 45° relative inclination, LSFM image stacks had to be processed to obtain isotropic volumetric images properly aligned with the sample holder. Therefore, raw stacks underwent a composite affine transformation (scaling, shearing, and rotation) by means of a custom Python tool which generated images oriented according to the reference system of the laboratory, characterized by an isotropic voxel size of 3.64 μm (i.e., the original voxel depth produced by the stage step motion).

Adjacent, overlapping TPFM and LSFM image stacks were then aligned and fused using the open-source ZetaStitcher tool (https://github.com/lens-biophotonics/ZetaStitcher) [33]. ZetaStitcher efficiently identifies the optimal alignment of adjacent image tiles by evaluating the FFT-based cross-correlation of overlapping regions at selected depths, enabling the high-throughput stitching of large volumetric microscopy datasets. The nominal lateral tile overlaps of ≈50 and ≈100 μm, respectively adopted for adjacent image stacks acquired with the custom TPFM and LSFM setups, guaranteed that no spurious doubling of corresponding tissue structures took place within overlapping areas.

### Microscopy image segmentation

#### TPFM pixel classification

ilastik’s pixel classification workflow was used to generate 3D semantic segmentations of the TPFM image reconstructions, associating each voxel with predefined semantic class labels. Four separate classes were adopted: pRPS6-neurons, pRPS6+ neurons, blood vessels, and background. In detail, cell bodies expressing only the NeuN protein, thus solely associated with red fluorescence emission around 655 nm, were labeled as pRPS6-, whereas the pRPS6+ class was adopted when labeling neurons emitting light around 570 nm as well. pRPS6- and pRPS6+ cell detections were therefore mutually exclusive. ilastik’s random forest classifier, once trained by expert human operators, estimated the probability of each voxel belonging to each of the above classes, based on a set of multiscale voxel-wise features related to RGB intensity, edgeness, and texture. The adopted features more specifically comprised the output of Gaussian smoothing filters of varying full width at half maximum (multiscale intensity), the output of a range of Laplacian of Gaussian kernels (multiscale edgeness), and, finally, the eigenvalues of scaled Hessian matrices of second-order partial derivatives (multiscale texture). The spatial scales of interest were thoroughly selected according to the expected size of neuronal bodies. In detail, the feature scale space was sampled assuming minimum and maximum soma diameters of 15 and 25 μm, respectively, with a step size of 5 μm. The corresponding scales σ (i.e., the standard deviation of the smoothing filters performing the image scale selection) were then derived as follows:

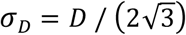

where *σ*_*D*_ is the optimal scale (in μm) where the local response to the presence of a binary sphere of diameter *D* is maximized (Lindeberg et al., 1992). Scale values in pixels, to be specified within ilastik’s feature selection widget, were finally obtained by taking into account the anisotropic voxel size of 1.21 μm × 1.21 μm × 2 μm (i.e., dividing by its lateral and longitudinal sizes).

The training of the classifier was conducted by three different operators on a random subset of TPFM image stacks, sampling and manually annotating one stack from each tiled reconstruction composing our dataset.

#### Neuronal body classification

pRPS6- and pRPS6+ cell probability maps returned by pixel classification were separately fed to the second object classification workflow and initially binarized by hysteresis thresholding. No preliminary smoothing was performed. In contrast to standard thresholding, this approach allowed on one hand to suppress noisy pixels within the background and, on the other hand, to preserve the and low thresholds of 80% and 50% were imposed on the input probability maps. Connected - component analysis [47] was then applied to label connected pixel regions belonging to the foreground class and detect individual 3D objects.

Object classification through object-level features, following the binarization of the pixel-wise semantic probability maps, aimed at preventing the inclusion of spurious neuronal bodies and, secondly, at addressing the incorrect merging of densely packed cells into a single object. Six properties were considered in this regard, namely: size of the segmented object, radii of the object (i.e., the eigenvalues returned by the principal components analysis of the object pixels’ coordinates), mean object intensity, intensity variance, convex hull volume, and object convexity (defined as the ratio between the object volume and the one of its convex hull). The number of object features was purposely kept low to avoid overfitting regimes leading to poor generalization capabilities, as the size of the training set coincided with the number of annotated cells (significantly lower than the number of pixels annotated in the previous workflow). Again, the object classifier training was performed by three separate operators. Once trained, this classifier enabled us to automatically discriminate single cells, merged objects (with the number of fused cell bodies), and likely false detections.

#### Quantitative cell population analysis

Quantitative data returned by ilastik’s object segmentation workflow were further processed to assess the specific spatial distribution and morphology of pRPS6- and pRPS6+ cells. Four properties were investigated, namely: cell volume, cell ellipticity, local cell density, and clustering degree.

Cell volume was directly provided by ilastik (being one of the features employed for object classification), whereas cell ellipticity was estimated as the ratio between the shortest and longest radii of each segmented object. The subsequent statistical analysis of these two features only focused on single-cell detections, in order to avoid any bias due to merged neuronal bodies.

On the other hand, local cell density and spatial clustering were evaluated as previously described (Silvestri et al., 2021). Specifically, the spatial clustering of neurons was evaluated using the 3D Ripley K-function (Ripley, 1976):

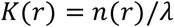

where *n*(*r*) represents the number of neighboring cells within a radius *r*, and *λ* is the local cell density, evaluated in a sphere of diameter 300 μm surrounding the centroid of each neuronal body. This reference volume is indeed substantially larger than the average cell but still smaller than the size of the analyzed TPFM image reconstructions. Under the hypothesis of a completely random spatial distribution, the expected value of *K*(*r*) corresponds to the sphere volume:

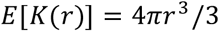

The basic principle of the formulated 3D cell clustering index, *I*, is that local violations of such a random condition can be quantified for each neuron by integrating the deviations from the above-expected value over a specific range of distances:

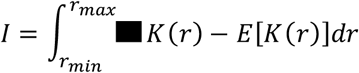

In the present work, a minimum and maximum radius of *r*_*min*_ = 10 μm (the approximate average soma radius), and *r*_*max*_ = 100 μm (below the radius of the reference sphere, used for local density estimation) were respectively selected, along with a discrete integration step of 10 μm.

Local cell density and spatial clustering of each segmented cell object were concurrently computed by means of an optimized Python tool leveraging multiprocessing and SciPy’s k-d tree implementation for fast nearest-neighbour lookup [31] to shrink the computation time required to handle our dataset of cell body detections. Regarding the spatial mapping of these two neuronal distribution properties, merged cell objects were carefully taken into consideration by transforming their original centroid into a proper number of random coordinates surrounding it (e.g., two 3D coordinates for an object classified as being formed by two cell bodies). Such coordinates were specifically obtained by first defining a set of random unit vectors, and then multiplying their length by distance values sampled from a uniform random distribution, bounded by the diameters corresponding to the 5% and 95% percentiles of the volume of single-body detections (18.9 μm and 21.2 μm, respectively, assuming a spherical shape). This mechanism aimed at preventing the underestimation of local density (and thus clustering) values, which would be understandably significant if the presence of these merged objects, corresponding in fact to tightly packed cell bodies, were neglected.

### Statistical analysis

To ascertain if the aforementioned neuronal cell measures were subject-specific, that is, dependent on the pertinent diseases, or influenced by the presence of phosphorylated S6 ribosomal proteins, data were initially assessed using the nonparametric Scheirer-Ray-Hare test (Scheirer et al., 1976). This was carried out by means of a custom R script (R Core Team, 2021). Subsequently, nonparametric Mann-Whitney U tests for independent samples were performed for pairwise comparisons across subjects and channels (i.e., pRPS6-versus pRPS6+ neurons). Pairwise Mann-Whitney tests were implemented through an original Python script, using SciPy’s stats module. All results were deemed statistically significant if their corresponding p-value was below 0.05. Bonferroni correction was applied in the case of the pairwise Mann-Whitney U tests, adjusting the above significance threshold in relation to the overall number of possible comparisons (i.e., *⍺*/6 ≈ 0.008). Dissimilarity between the compared data distribution pairs, *p*(*x*) and *q*(*x*), was described in terms of their Hellinger distance, *H*(*p*,*q*) | *0* ≤ *H*(*p*,*q*) ≤ *1* (Hellinger, 1909). For a pair of discrete probability distributions, *p*_*d*_ = (*p_1_*, …, *p*_*n*_) and *q*_*d*_ = (*q_1_*, …, *q*_*n*_), this metric can be computed as:

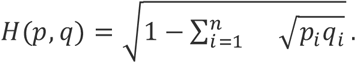

## Acknowledgements

We thank Dr. Giuseppe de Vito (from University of Florence, National Research Council – National Institute of Optics, and the European Laboratory for Non-Linear Spectroscopy, LENS) for his useful suggestions on the adopted statistical analysis approaches. We express our gratitude to the donors involved in the body and tissue donation programs who made this study possible by generously donating their brains to science.

## Author contributions statement

DDM and IC optimized the deparaffinization protocol with input from AE. DDM and JR tested the compatibility with immunolabeling and prepared the sample. DDM, FC, and LP performed light-sheet imaging. IC performed two-photon imaging. MS implemented the ML-based image segmentation pipeline, developed the tools for the investigation of quantitative cytoarchitectural features, and carried out the statistical analysis. MS, SB, and GM performed the microscopy image postprocessing for the generation of the 3D tissue sample reconstructions. MS, DDM, and BL performed the manual image annotation for training the ilastik pixel and object classifiers. AE, AP, RD, RG, DB, CP, VC, and RG provided the samples and helped in interpreting the results. RG and DB performed classical staining of the hippocampus sample. CP, VC, and RG performed the genetic analysis of pediatric samples. FSP acquired the financial support and provided the equipment employed in this work. IC conceived and supervised the study. DDM, MS, and IC interpreted the results and wrote the manuscript with inputs from all the authors.

## Supplementary figures and tables

**Figure S1.**
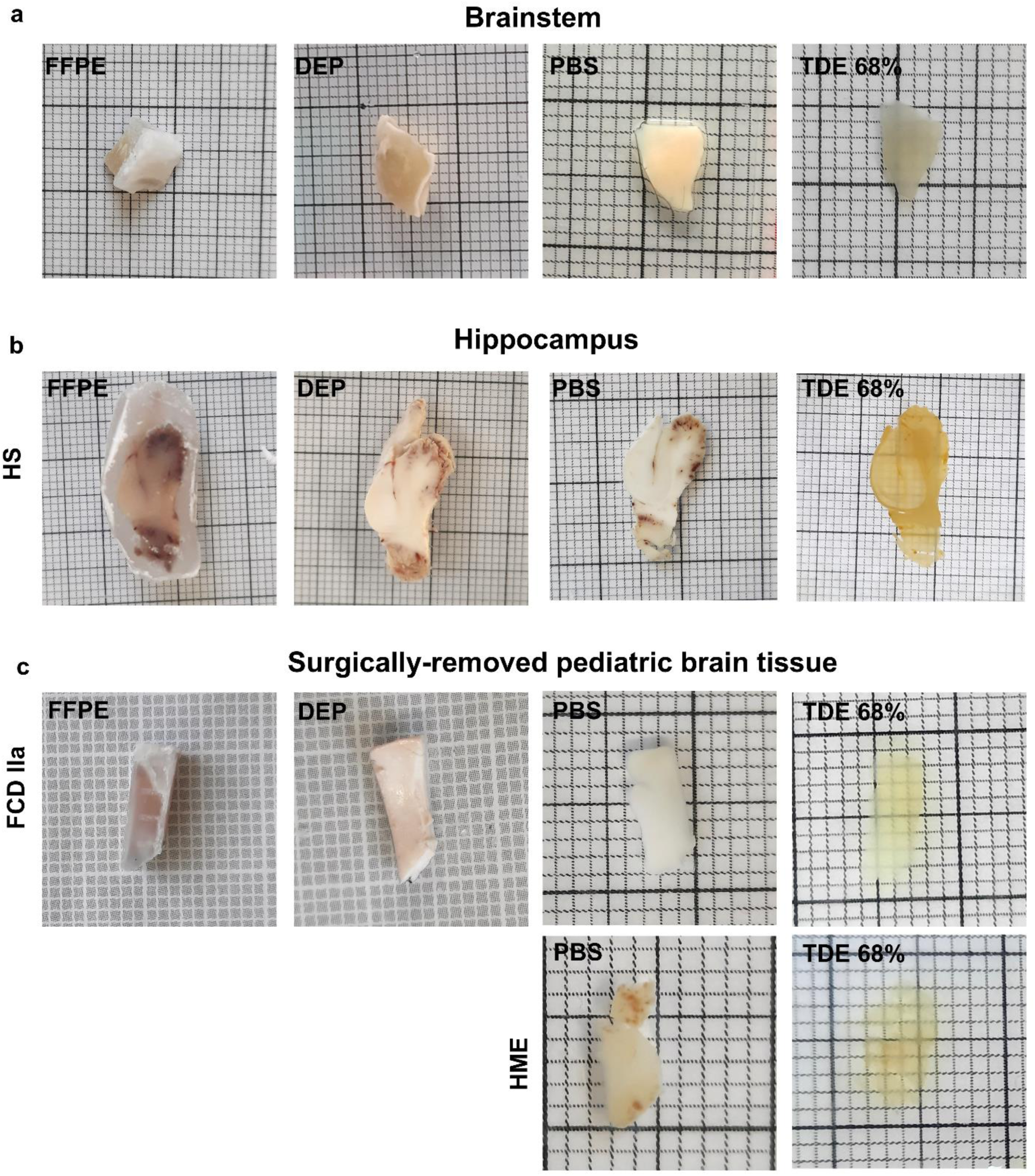
Brain tissue throughout the whole pipeline. Images showing the different specimens before the whole process (FFPE, Formalin-Fixed Paraffin-Embedded), after the deparaffinization step (DEP), single 500 µm-thick slices before SHORT (PBS), and after clearing, labeling and RI matching (TDE 68%). **a)** Postmortem adult human brainstem. **b)** Postsurgical hippocampus from a patient with hippocampal sclerosis (hippocampus HS). **c)** Brain specimens surgically removed from pediatric patients with focal cortical dysplasia type IIa (FCDIIa) and hemimegalencephaly (HME).

**Figure S2.**
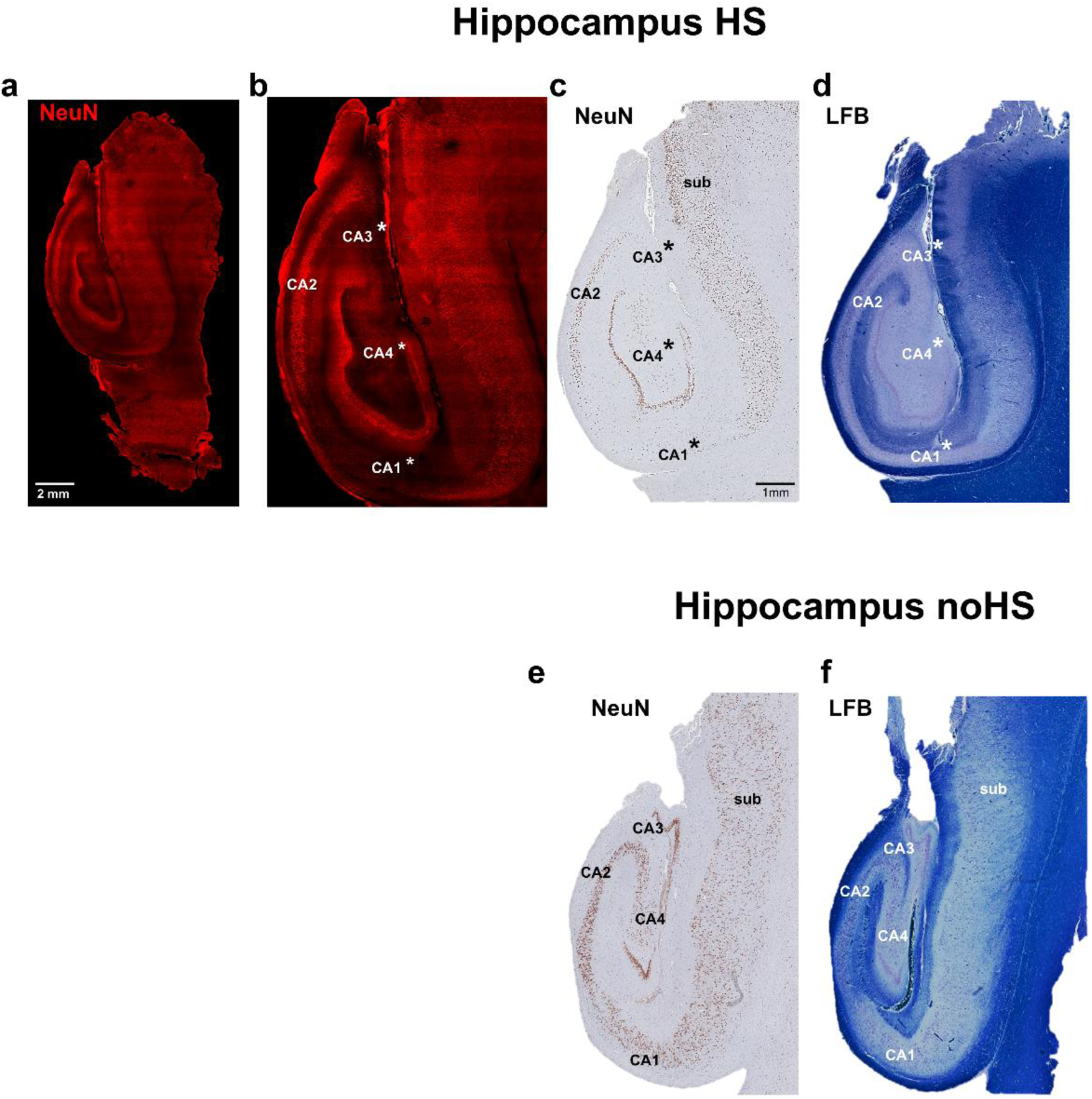
LSFM reconstruction and histological staining on human hippocampus. **a, b)** Maximum intensity projection images showing a mesoscopic reconstruction of postsurgical human hippocampus from a patient with hippocampal sclerosis (HS), shown in Figure 2e, labeled for NeuN (red). Note the prominent pyramidal cell loss in CA1, CA3 and CA4 (indicated by asterisks) typical sign of hippocampal sclerosis. **c, d)** NeuN-immunostaining (c) and Luxol Fast Blue-staining (d, LFB) on FFPE sections, obtained from the same tissue block prior to the cleaning protocol, confirm this pattern of neuronal loss. **e, f)** For comparison, an example of NeuN-immunostaining (e) and Luxol Fast Blue-staining (f, LFB) on FFPE tissue section from a noHS specimen is shown to highlight the normal cellular density in the different CA sectors.

**Figure S3.**
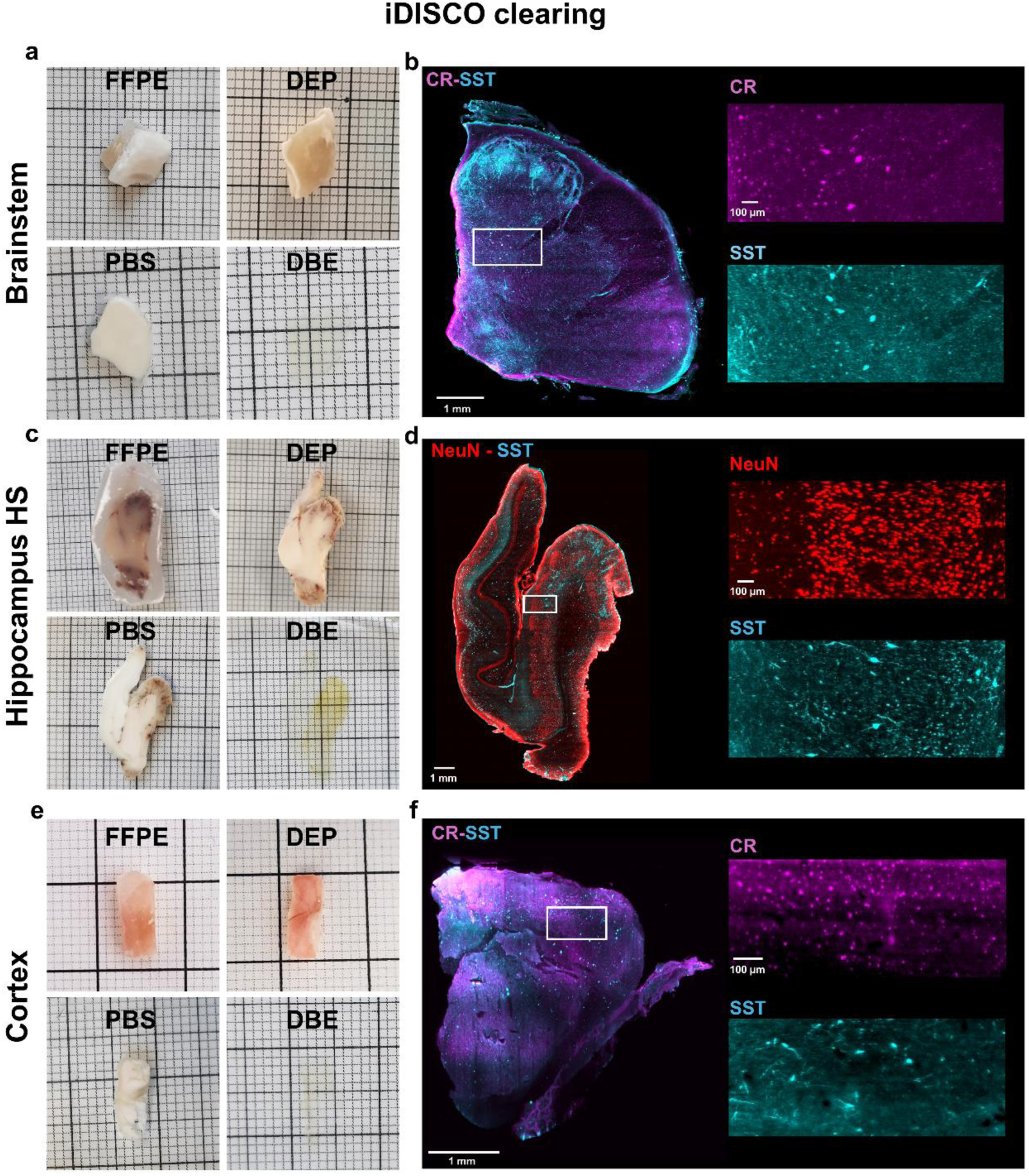
Volumetric reconstruction of deparaffinized human brain slabs (500 μm in thickness) processed with iDISCO. **a)** Images showing the postmortem adult human brainstem before the whole process (FFPE), after the deparaffinization step (DEP), single 500 µm-thick slices before iDISCO (PBS) and after clearing, labeling, and RI matching (DBE). **b)** Maximum intensity projection image (3.64 µm isotropic resolution) showing a mesoscopic reconstruction of (a) labeled for Calretinin (CR, magenta) and Somatostatin (SST, cyan). Scale bar: 1 mm. On the right are magnified insets showing the single marker used. Scale bar: 100 µm. **c)** Images showing the postsurgical adult human hippocampus from a patient with hippocampal sclerosis before the whole process (FFPE), after the deparaffinization step (DEP), single 500 µm-thick slices before iDISCO (PBS) and after clearing, labeling and RI matching (DBE). **d)** Maximum intensity projection image showing a mesoscopic reconstruction of (c) labeled for NeuN (red) and Somatostatin (SST, cyan). Scale bar: 1 mm. On the right are magnified insets showing the single marker used. Scale bar: 100 µm. **e)** Images showing a surgically removed specimen from a pediatric brain before the whole process (FFPE), after the deparaffinization step (DEP), single 500 µm-thick slices before iDISCO (PBS) and after clearing, labeling and RI matching (DBE). **f)** Maximum intensity projection image showing a mesoscopic reconstruction of (e) labeled for Calretinin (CR, magenta) and Somatostatin (SST, cyan). Scale bar:

**Table S1:**

| Antibodies | Company | Cat. n. | Dilution |
| --- | --- | --- | --- |
| Anti-NeuN Antibody | Merck Life Science S.R.L. | ABN91 | 1:50 |
| Phospho-S6 ribosomal protein, Ser240/244 | Cell Signaling | 5364 | 1:100 |
| Calretinin Polyclonal antibody | ProteinTech | 12278_1_AP | 1:200 |
| Somatostatin Antibody YC7 | Santa Cruz Biotechnology | sc-47706 | 1:200 |
| Goat Anti-Chicken IgY H&L (Alexa Fluor® 647) | Abcam | ab150171 | 1:200 |
| Goat Anti-Rabbit IgG H&L (Alexa Fluor® 568) | Abcam | 175471 | 1:200 |
| Recombinant Alexa Fluor® 488 Anti-GFAP | Abcam | ab194324 | 1:200 |
| Donkey Anti-Rat IgG H&L (Alexa Fluor® 568) | Abcam | ab175475 | 1:200 |
| DAPI (Dilactate) | Thermo Fisher Scientific | D3571 | 1:100 |

**Table S2:**

| Subject | pRPS6+ [%] | mean pRPS6+ density [cell / mm<sup>3</sup>] | mean total density [cell / mm<sup>3</sup>] | tissue volume [mm<sup>3</sup>] |
| --- | --- | --- | --- | --- |
| S<sub>1</sub> | 3.1 | 344 | 11257 | 6.9 |
| S<sub>2</sub> | 35.4 | 3230 | 9123 | 4.6 |

**Table S3:**
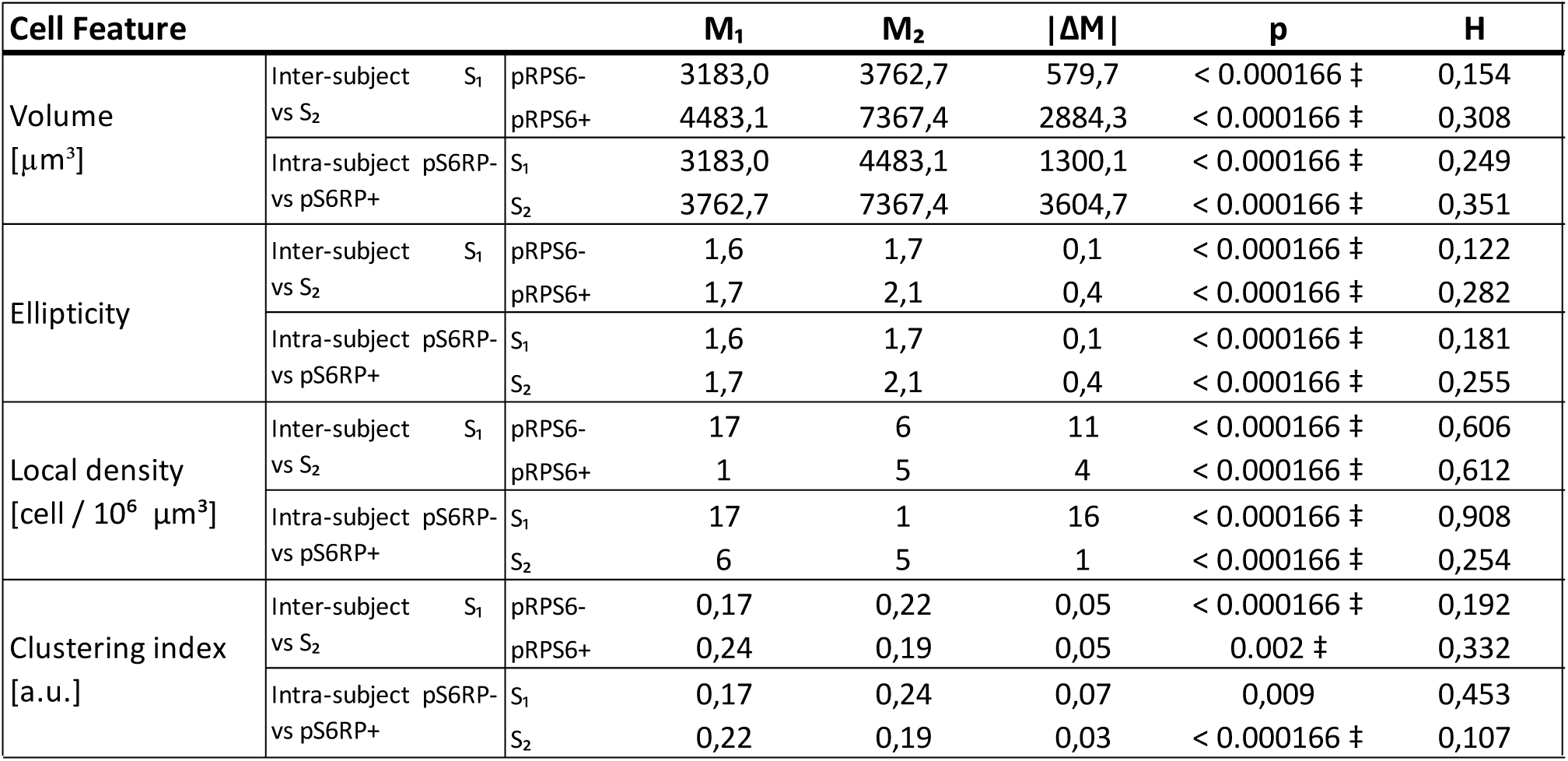

**Table S4:**

| Samples | Clearing days | Primary antibodies incubation days | Secondary antibodies incubation days |
| --- | --- | --- | --- |
| S<sub>1</sub> (HME) | 3 | 2 | 1 |
| S<sub>2</sub> (FCDIIa) | 3 | 2 | 1 |
| Brainstem | 4 | 3 | 2 |
| Hippocampus | 6 | 6 | 5 |

